# Projection-specific filtering of delta-theta input by layer 5 neurons in mouse secondary visual cortex

**DOI:** 10.64898/2026.11.06.763938

**Authors:** Hugh Takemoto, Zhiwen Ye, Ede Rancz

**Affiliations:** INSERM, INMED, Aix Marseille Univ, Marseille, France; The Francis Crick Institute, London, United Kingdom; Aix Marseille Univ, Turing Centre for Living systems, Marseille, France

**Author notes:** Institute of Neuromodulation and Cognition, Shenzhen Medical Academy of Research and Translation (SMART), Shenzhen, China (Zhiwen Ye).

## Abstract

Delta-and theta-band oscillations are thought to coordinate cortical activity and to convey top-down signals, but how the different output neurons of a cortical area transform such rhythmic input is less well understood. Layer 5 contains two projection classes, extratelencephalic (ET) and intratelencephalic (IT) neurons, which carry different information to different targets. Here we asked whether they also differ in how they process slow rhythmic input in the delta–theta range. We combined whole-cell recordings in three stimulus regimes in mouse medial secondary visual cortex (V2M) with local field potential (LFP) recordings in awake mice. With small inputs, IT cells summated strongly and acted as low-pass filters, whereas ET cells resonated at low frequencies, forming moderate-resonance (ET1, about 3 Hz) and high-resonance (ET2, about 5 Hz) subgroups. With large inputs, ET cells amplified depolarisation more than hyperpolarisation, at similar resonance frequencies. When driven to fire by a 4 Hz sinusoid at threshold, cells of all classes fired at a similar phase and kept it nearly constant across cycles. In awake mice, visual stimulation increased deep-layer LFP power across about 1–6 Hz, a band that spans the mean ET1 and ET2 resonance frequencies. Thus, the two output classes of layer 5 in V2M filter slow rhythmic input differently, but once firing, they show no detectable difference in spike timing.

## Introduction

Oscillations in the local field potential (LFP) are ubiquitous in cortical structures and occur across a wide range of spatial and temporal scales. Rhythms in the theta band have been proposed to synchronise activity within and between cortical areas, to convey top-down signals such as expectation, and to support the integration of information across modalities (Buzsáki, 2002; Colgin, 2013; Fournier et al., 2020). Delta-band (about 1-4 Hz) activity in the awake cortex has been proposed to serve sensory selection by phase-locking to the temporal structure of attended stimuli (Schroeder and Lakatos, 2009), and in the somatosensory cortex of awake mice it is linked to respiration (Ito et al., 2014). Detailed biophysical models of cortical circuits suggest that pyramidal neurons are the principal contributors to the rhythms measured extracellularly (Dura-Bernal et al., 2023). If such rhythmic activity carries information, the neurons embedded in it must be able to read it out. How individual neurons convert rhythmic input into spike output is therefore central to understanding what such rhythms can convey. Here we focus on the delta–theta range (about 1-8 Hz), which contains both the cellular resonances and the visually evoked rhythm studied below.

Higher-order visual areas are a natural place to ask this question, as they are likely to receive a broader diversity of long-range input than primary sensory cortex. In the medial secondary visual cortex (V2M), rabies tracing from ET neurons labels many presynaptic neurons in the retrosplenial cortex (RSP), which, together with V2M itself and primary visual cortex, contributes most of their input cells (Galloni et al., 2022). This input is not confined to the apical tuft: subcellular optogenetic mapping shows that RSP axons target the apical tuft, oblique and basal dendrites to a similar extent, with the peak input on perisomatic dendrites in most neurons (Galloni et al., 2022). A rhythmic RSP signal in this range would therefore reach perisomatic as well as distal dendrites, making somatic integration relevant for this input. Widefield voltage imaging has shown that visually evoked and spontaneous 5 Hz oscillations in V1 are accompanied by similar oscillations in RSP (White et al., 2026), consistent with this anatomical route. Whether such an oscillation is present in V2M itself, and specifically in layer 5 where the output neurons reside, has not been tested.

Layer 5 is the principal output layer of the cortical column. Its excitatory population divides into cells projecting to the superior colliculus and brainstem (extratelencephalic, ET) and cells projecting to ipsilateral cortex or striatum (intratelencephalic, IT). These populations are functionally as well as anatomically distinct. In visual cortex, they respond differently to a range of visual stimuli (Lur et al., 2016), and ET cells, but not their IT neighbours, encode sensory and motor task information and are selectively required for performance in a visually guided task (Tang and Higley, 2020).

Differences in intrinsic properties may contribute to this distinction. In rat medial prefrontal cortex, ET cells act as band-pass filters resonating at 3–6 Hz and summate EPSP trains weakly, whereas IT cells act as low-pass filters and summate strongly (Dembrow et al., 2010). Resonance of this kind arises when a slow, voltage-dependent conductance opposes changes in membrane potential, acting as a phenomenological inductance: the hyperpolarisation-activated current Ih at membrane potentials around and below rest, and the M-current (Kv7/KCNQ) at more depolarised Vm (Hutcheon and Yarom, 2000; Mishra and Narayanan, 2025). Modelling of L5 ET neurons indicates that Ih, together with a shunting conductance, shapes resonance frequency, impedance magnitude and phase (Kelley et al., 2021), and that these cells should respond with different phase to depolarising and hyperpolarising drive once inputs become large enough to engage voltage-gated channels asymmetrically (Kelley et al., 2023). However, two questions remain open. First, resonance is conventionally measured with small inputs, which keep voltage-dependent conductances close to their resting state and therefore evoke approximately linear responses, whereas physiological inputs are often large enough to recruit these conductances and evoke non-linear responses, so the linear impedance profile may not predict the response to such input. Second, resonance is a subthreshold property. Whether it carries over to spike timing, and whether that timing remains stable across successive cycles of a sustained oscillation, as it need not in models of L5 neurons (Kelley et al., 2023), has not been examined in identified L5 projection classes. The phase at which a neuron fires relative to an ongoing rhythm is itself a coding dimension (Lisman and Jensen, 2013), and phase coding requires spike phase to be stable over a bout.

Here, we characterised the responses of V2M layer 5 neurons in three stimulus regimes: small subthreshold inputs (linear regime), large subthreshold inputs (non-linear regime) and suprathreshold drive. We then asked whether a rhythm in this frequency range is present in the deep layers of V2M in vivo. With small inputs, IT cells summated strongly and acted as low-pass filters, whereas ET cells resonated, forming moderate-resonance (ET1, about 3 Hz) and high-resonance (ET2, about 5 Hz) subgroups. With large inputs, ET cells amplified depolarisation more than hyperpolarisation. When driven to fire at 4 Hz, cells of all classes fired at a similar phase and kept it nearly constant across cycles. In vivo, visual stimulation increased deep-layer LFP power in V2M across about 1–6 Hz, a band that did not follow the stimulus frequency and spans the mean ET1 and ET2 resonance frequencies. As ET cells form the output of the cortical column and provide little local excitatory output (Harris and Shepherd, 2015), they are unlikely to pace this rhythm. Our results show that the two output channels of layer 5 filter slow rhythmic input differently, with no detectable difference in spike timing once they fire.

## Methods

### Animals

For in vitro experiments, male wild type (C57BL/6J) and transgenic (Colgalt2-Cre-NF107::Ai14 and Tlx3-Cre::Ai14) mice were used in all experiments. The Colgalt2-Cre::Ai14 mice expressed fluorescent protein td-tomato in ET neurons in V2M. The Tlx3-Cre::Ai14 mice expressed td-tomato in L5 IT neurons in V2M. In vitro animal experiments were prospectively approved by the Animal Care and Use Committee of the Institut National de la Santé et de la Recherche Médicale (INSERM) and authorised by the Ministère de l’Education Nationale de l’Enseignement Supérieur et de la Recherche after evaluation by a local ethics committee (agreement number APAFIS #38603-2022091917383392).

For in vivo experiments, recordings were obtained from five adult mice (both sexes, aged between 2-6 months). In vivo animal experiments were prospectively approved by the local ethics panel of the Francis Crick Institute and the UK Home Office under the Animals (Scientific Procedures) Act 1986 (PPL: 70/8935). All surgeries were performed under isoflurane anaesthesia, and every effort was made to minimise suffering.

### In vitro slice preparation

Young adult mice (5–12 weeks old) were deeply anaesthetised with isoflurane before they were decapitated. Brains were removed and placed in oxygenated ice-cold slicing solution containing (in mmol/L): 110 choline chloride, 2.2 potassium chloride, 1.09 NaH2PO4, 6.12 MgCl2, 0.437 CaCl2, 23.8 NaHCO3, 11.1 glucose (osmolarity 305-310). Slices were cut coronally with a vibrating blade microtome (Leica VT1200S or Campden 7000smz-2) to 250 µm thickness at an angle of approximately 10 degrees to preserve apical dendrites. Only slices containing the visual cortex were preserved (judged to be between 2.3 and 4.3 mm posterior to bregma). Slices were maintained in a submerged chamber containing artificial cerebrospinal fluid (ACSF) containing the following (in mM): NaCl 125, KCl 2.5, NaHCO3 26, NaH2PO4.H2O 1.25, Glucose 25, CaCl2 1, MgCl2 2 (osmolarity 305-310). For the first 30-60 minutes of incubation slices were maintained at body temperature 35 Celsius. Afterwards slices were kept at room temperature 22 degrees, all solutions and chambers were bubbled with carbogen (95% O2 / 5% CO2).

### In vitro whole-cell recording

After the incubation period slices were individually transferred to the recording chamber which were perfused with ACSF at a rate of 6mL/ min. The ACSF was heated to body temperature (35 Celsius) with an inline heater for all experiments. Synaptic blockers were added to the ACSF to block glutamatergic (kynurenic acid, 1mM) and GABAergic (picrotoxin, 50µM) synaptic input.

Filamented borosilicate thick-walled glass (GC150F, Phymep, France) micropipettes were pulled to obtain an electrode resistance of 3-6MΩ. The electrodes were filled with intracellular solution optimised for current clamp recording containing the following (in mM): KMeSO4 130, KCl 5, HEPES 10, EGTA 1, CaCl2 0.1, Mg2-ATP 4, Na-GTP 0.4, sodium phosphocreatine 5, 0.5% (w/v) biocytin hydrochloride, osmolarity 290-295 mOsm; pH adjusted to 7.3 with KOH.

Recordings were performed using a Scientifica SliceScope Pro 3000 microscope equipped with a 40x/0.8NA objective and an infrared (IR) Dodt Gradient Contrast system. Electrodes and recording headstages were mounted on motorised micromanipulators (Scientifica MicroStar). Electrode offset was set above the slice and positive pressure was applied to the pipette before lowering it into the slice. An oscilloscope was used to measure the resistance of the pipette tip during repetitive - 10mV voltage steps. Once the pipette was above the surface of the target cell the positive pressure was released and gentle suction was applied until a gigaohm seal was formed. Whole cell configuration was achieved by short suction pulses. Most cells had an access resistance of 10–25 MΩ. Cells were excluded from the analysis if the access resistance exceeded 50 MΩ or the holding current exceeded 100 pA during the recording.

Cells were recorded with a resting membrane potential of about −70 mV using small current injections if necessary. Voltage measurements were not corrected for the junction potential.

Linear resonance properties were measured with a ±20 pA, 30 s chirp current whose frequency increased exponentially from 0.5 to 50 Hz. For non-linear resonance properties, we used a ±80 pA, 30 s chirp current whose frequency increased linearly from 0.2 to 20 Hz.

To quantify summation, we injected five αEPSC waveforms (amplitude 20 pA) at 40 Hz and defined the summation ratio as the ratio between the peak voltage at the 5th and 1st current injection. To measure sag, we injected a 500 ms hyperpolarising current step and calculated the sag percentage (the magnitude of the sag with respect to the peak hyperpolarisation and the steady state, measured over the last 50 ms of the step). The step amplitude was adjusted so that the initial hyperpolarising peak reached about −100 mV. Action potential threshold was measured from an average spike waveform obtained from a 2x rheobase voltage trace. Spikes occurring 150–400 ms after step onset (to exclude bursts) were extracted and averaged. The first and third derivatives of the average spike waveform were computed. Threshold was defined as the membrane potential at which dV/dt first reached 10% of its maximum during the spike upstroke. This value was also the reference for AHP amplitude and latency. A second estimate, the potential at the peak of d3V/dt3, was computed for comparison. Threshold detection was verified visually for every cell. Adaptation ratio was measured from the spikes in the 2x rheobase voltage trace. Spikes were counted in the first half (0–250 ms) and second half (250–500 ms) of each trace. The mean spike count per half was taken across all repetitions. Adaptation ratio was calculated as 1 − (mean spikes in second half ÷ mean spikes in first half). A value of 0 means no spike-frequency adaptation (equal firing in both halves), and values approaching 1 mean strong adaptation (firing largely ceases in the second half). A cell with no spikes detected during the step was assigned a ratio of 1 (full depolarisation block).

### In vitro statistics and analysis

All statistics were performed using Python. Data normality was assessed using Shapiro–Wilk tests. Independent t-tests or Mann–Whitney U tests were used to compare two groups; one-way ANOVA with Tukey’s post hoc test, or the Kruskal–Wallis H test with Dunn’s post hoc tests (Bonferroni-corrected), was used for three groups. Peak and trough measures from the same cell were compared with paired t-tests, Bonferroni-corrected across the four measures. Values are mean ± SEM unless stated otherwise; all comparisons are listed in Table 1.

**Table 1.**
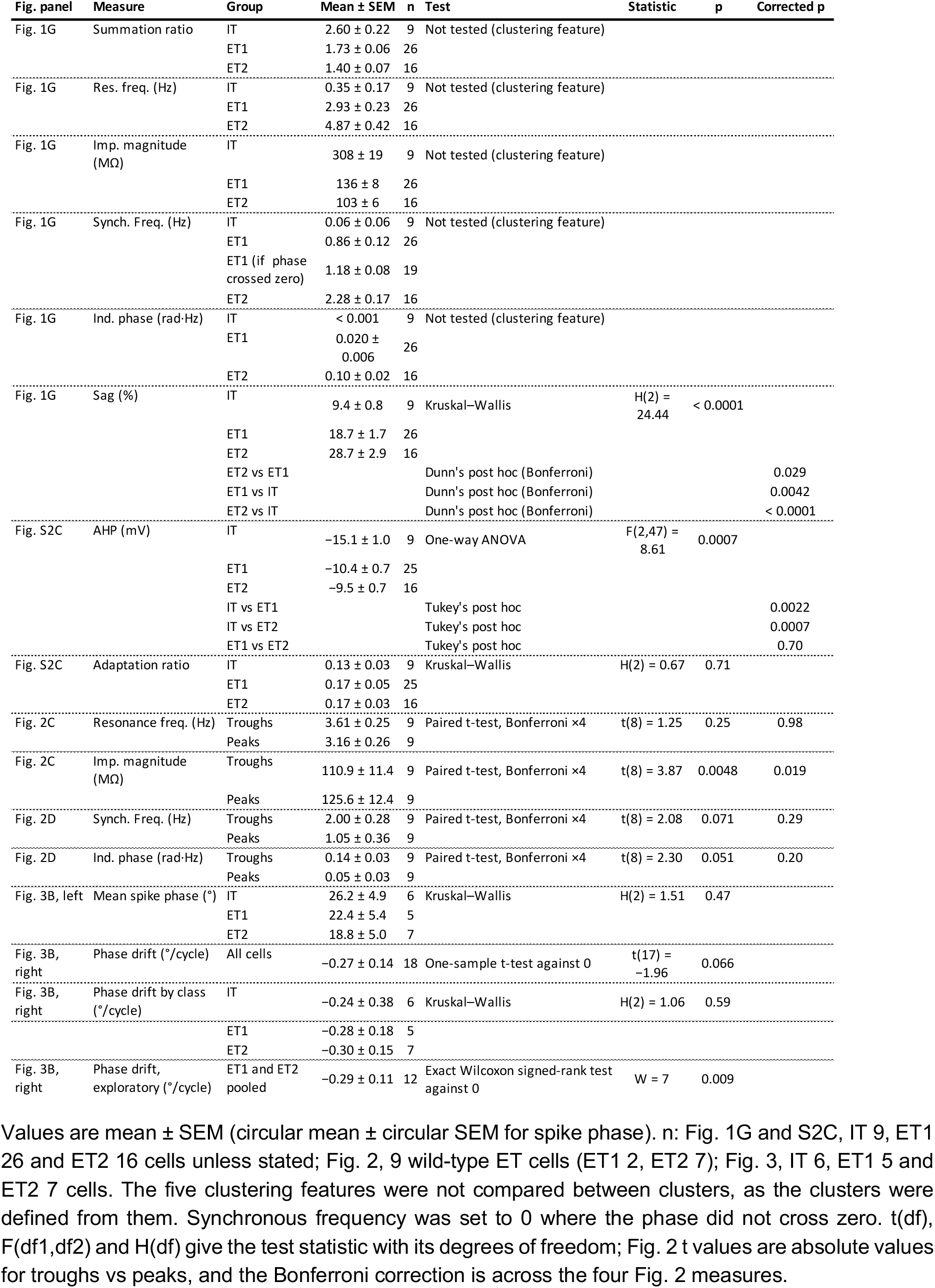
Group values and statistical comparisons.

| Fig. panel | Measure | Group | Mean ± SEM | n | Test | Statistic | p | Corrected p |
|---|---|---|---|---|---|---|---|---|
| Fig. 1G | Summation ratio | IT | 2.60 ± 0.22 | 9 | Not tested (clustering feature) |  |  |  |
|  |  | ET1 | 1.73 ± 0.06 | 26 |  |  |  |  |
|  |  | ET2 | 1.40 ± 0.07 | 16 |  |  |  |  |
| Fig. 1G | Res. freq. (Hz) | IT | 0.35 ± 0.17 | 9 | Not tested (clustering feature) |  |  |  |
|  |  | ET1 | 2.93 ± 0.23 | 26 |  |  |  |  |
|  |  | ET2 | 4.87 ± 0.42 | 16 |  |  |  |  |
| Fig. 1G | Imp. magnitude (MΩ) | IT | 308 ± 19 | 9 | Not tested (clustering feature) |  |  |  |
|  |  | ET1 | 136 ± 8 | 26 |  |  |  |  |
|  |  | ET2 | 103 ± 6 | 16 |  |  |  |  |
| Fig. 1G | Synch. Freq. (Hz) | IT | 0.06 ± 0.06 | 9 | Not tested (clustering feature) |  |  |  |
|  |  | ET1 | 0.86 ± 0.12 | 26 |  |  |  |  |
|  |  | ET1 (if phase crossed zero) | 1.18 ± 0.08 | 19 |  |  |  |  |
|  |  | ET2 | 2.28 ± 0.17 | 16 |  |  |  |  |
| Fig. 1G | Ind. phase (rad·Hz) | IT | < 0.001 | 9 | Not tested (clustering feature) |  |  |  |
|  |  | ET1 | 0.020 ± 0.006 | 26 |  |  |  |  |
|  |  | ET2 | 0.10 ± 0.02 | 16 |  |  |  |  |
| Fig. 1G | Sag (%) | IT | 9.4 ± 0.8 | 9 | Kruskal–Wallis | H(2) = 24.44 | < 0.0001 |  |
|  |  | ET1 | 18.7 ± 1.7 | 26 |  |  |  |  |
|  |  | ET2 | 28.7 ± 2.9 | 16 |  |  |  |  |
|  |  | ET2 vs ET1 |  |  | Dunn's post hoc (Bonferroni) |  |  | 0.029 |
|  |  | ET1 vs IT |  |  | Dunn's post hoc (Bonferroni) |  |  | 0.0042 |
|  |  | ET2 vs IT |  |  | Dunn's post hoc (Bonferroni) |  |  | < 0.0001 |
| Fig. S2C | AHP (mV) | IT | −15.1 ± 1.0 | 9 | One-way ANOVA | F(2,47) = 8.61 | 0.0007 |  |
|  |  | ET1 | −10.4 ± 0.7 | 25 |  |  |  |  |
|  |  | ET2 | −9.5 ± 0.7 | 16 |  |  |  |  |
|  |  | IT vs ET1 |  |  | Tukey's post hoc |  |  | 0.0022 |
|  |  | IT vs ET2 |  |  | Tukey's post hoc |  |  | 0.0007 |
|  |  | ET1 vs ET2 |  |  | Tukey's post hoc |  |  | 0.70 |
| Fig. S2C | Adaptation ratio | IT | 0.13 ± 0.03 | 9 | Kruskal–Wallis | H(2) = 0.67 | 0.71 |  |
|  |  | ET1 | 0.17 ± 0.05 | 25 |  |  |  |  |
|  |  | ET2 | 0.17 ± 0.03 | 16 |  |  |  |  |
| Fig. 2C | Resonance freq. (Hz) | Troughs | 3.61 ± 0.25 | 9 | Paired t-test, Bonferroni ×4 | t(8) = 1.25 | 0.25 | 0.98 |
|  |  | Peaks | 3.16 ± 0.26 | 9 |  |  |  |  |
| Fig. 2C | Imp. magnitude (MΩ) | Troughs | 110.9 ± 11.4 | 9 | Paired t-test, Bonferroni ×4 | t(8) = 3.87 | 0.0048 | 0.019 |
|  |  | Peaks | 125.6 ± 12.4 | 9 |  |  |  |  |
| Fig. 2D | Synch. Freq. (Hz) | Troughs | 2.00 ± 0.28 | 9 | Paired t-test, Bonferroni ×4 | t(8) = 2.08 | 0.071 | 0.29 |
|  |  | Peaks | 1.05 ± 0.36 | 9 |  |  |  |  |
| Fig. 2D | Ind. phase (rad·Hz) | Troughs | 0.14 ± 0.03 | 9 | Paired t-test, Bonferroni ×4 | t(8) = 2.30 | 0.051 | 0.20 |
|  |  | Peaks | 0.05 ± 0.03 | 9 |  |  |  |  |
| Fig. 3B, left | Mean spike phase (°) | IT | 26.2 ± 4.9 | 6 | Kruskal–Wallis | H(2) = 1.51 | 0.47 |  |
|  |  | ET1 | 22.4 ± 5.4 | 5 |  |  |  |  |
|  |  | ET2 | 18.8 ± 5.0 | 7 |  |  |  |  |
| Fig. 3B, right | Phase drift (°/cycle) | All cells | −0.27 ± 0.14 | 18 | One-sample t-test against 0 | t(17) = −1.96 | 0.066 |  |
| Fig. 3B, right | Phase drift by class (°/cycle) | IT | −0.24 ± 0.38 | 6 | Kruskal–Wallis | H(2) = 1.06 | 0.59 |  |
|  |  | ET1 | −0.28 ± 0.18 | 5 |  |  |  |  |
|  |  | ET2 | −0.30 ± 0.15 | 7 |  |  |  |  |
| Fig. 3B, right | Phase drift, exploratory (°/cycle) | ET1 and ET2 pooled | −0.29 ± 0.11 | 12 | Exact Wilcoxon signed-rank test against 0 | W = 7 | 0.009 |  |

Values are mean ± SEM (circular mean ± circular SEM for spike phase). n: Fig. 1G and S2C, IT 9, ET1 26 and ET2 16 cells unless stated; Fig. 2, 9 wild-type ET cells (ET1 2, ET2 7); Fig. 3, IT 6, ET1 5 and ET2 7 cells. The five clustering features were not compared between clusters, as the clusters were defined from them. Synchronous frequency was set to 0 where the phase did not cross zero. t(df), F(df1,df2) and H(df) give the test statistic with its degrees of freedom; Fig. 2 t values are absolute values for troughs vs peaks, and the Bonferroni correction is across the four Fig. 2 measures.

PCA was performed on five features per cell (summation ratio, resonance frequency, resonance amplitude, synchronous frequency, and area of inductive phase), extracted for all cells with complete data across these five measures. Features were standardised (z-scored) using a StandardScaler fit across cells before dimensionality reduction; the first two principal components were retained for visualisation. K-means clustering was performed on the standardised five-feature data directly, with the number of clusters (k=3) chosen following inspection of silhouette and elbow plots. The reference clustering solution was obtained from a single k-means run with one centroid initialisation (n_init=1, random_state=45).

Cluster reliability was assessed separately by re-running k-means 101 times, each with a single initialisation (n_init=1) and a distinct random seed (0–100). Because k-means cluster labels are arbitrary across runs, each run’s labels were aligned to the reference solution’s label space using the Hungarian algorithm applied to a run-vs-reference contingency table, maximising label overlap before comparison. For each cell, a stability score was defined as the fraction of the 101 aligned runs in which the cell was assigned to its own modal (most frequently assigned) cluster label. Cells were split into “stable” (core) and “borderline” groups by a median split on this stability score.

### Calculating the linear impedance profile

We calculated the impedance profile using:

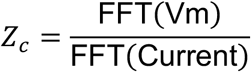

*Z_c_* is a complex valued function where FFT(Vm) is the fast Fourier transform of the voltage response and FFT(Current) is the fast Fourier transform of the injected current. From the impedance we extracted the real valued resistance (*R*) and the imaginary valued reactance (*X*). From *R* and *X* we calculated the impedance amplitude:

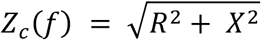

Resonance frequency was defined as the frequency at which *Z_c_* is maximal (this was defined as 0 for cells with no discernible bump in the impedance magnitude profile). We defined impedance phase (*Φ_c_*) as the temporal relationship between the input and the output:

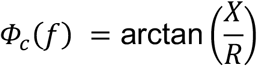

Synchronous frequency was defined as the frequency at which *Φ_c_*= 0 and peaks in Vm and the Current are synchronised. When *Φ_c_*> 0 the peaks in Vm precede peaks in the current. We refer to this as leading or inductive phase. Total inductive phase is defined as:

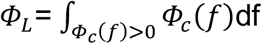

The area of the *Φ_c_* curve above zero. If there is no inductive phase and *Φ_c_* < 0 for all frequencies then we set *Φ_L_* and synchronous frequency to zero.

### Calculating non-linear resonance and phasic profiles

We injected a large (±80 pA) linearly increasing chirp (0.2 to 20 Hz) to measure the non-linear voltage responses of V2M L5 ETs to time-variant stimuli. Since the linear impedance analysis is insufficient for asymmetric non-linear responses, we used the analysis developed by Kelley et al. (2023) and Pena et al. (2019) to compute a non-linear analogue of impedance phase (+) and magnitude for hyperpolarisation and depolarisation. As in Kelley et al. (2023), we defined the upper (+) and lower (-) envelopes of the the Vm by finding the peaks (*V_n_^+^*) and troughs (*V_n_^-^*) for each stimulus cycle (n) and did the same for the Current (*I_n_^±^*). Then we defined the times of each peak and trough in Vm as *V_n_^±^* and the Current as *i_n_^±^*. The instantaneous phase of the Current was extracted using the Hilbert transform:

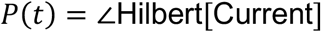

We defined our non-linear analogue for upper and lower envelopes of Vm (+/-) as:

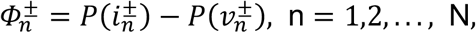

*N* represents the total number of cycles. We used a chirp that increased linearly over time therefore one can readily define *Φ*^±^(*f*). As with linear *Φ*, phase lead occurs when the voltage peaks/ troughs (*i_n_^±^*) led the current peaks/troughs (*i_n_^±^*) *Φ_n_^±^* > 0. Phase lag occurs, when the voltage peaks/ troughs come after current peaks/ troughs then *Φ_n_^±^* < 0. Synchronous frequency was defined as *Φ_n_^±^* = 0 when the peaks/ troughs of the voltage and current are in phase. We also calculated suprathreshold phasic responses by measuring *Φ*^+^of a single spike or the first spike in a burst.

As in Pena et al. (2019), we used the holding voltage (*V*_hold_) as a reference line to characterise any voltages above it to be positive and any voltages below it to be negative. For each frequency *f*, we measured the magnitude of the maximum (peaks *V*^+^) and minimum (troughs *V*^-^) voltage traces normalised to the amplitude of the input Current (*A*_in_). We refer to these two quantities, which depend on frequency (*f*) and have dimensions of impedance as upper impedance [*Z*^+^(*f*)] and lower impedance [*Z*^-^(*f*)]. Unlike the standard impedance, which averages the upper and lower responses, these measures reveal asymmetries between them.

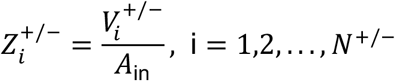

Where *V*^+/-^ is the absolute peak and trough distance from *V*_hold_ and *N*^+/-^ are the number of peaks and troughs in the time series. For each peak and trough, i, we check the corresponding frequency from the chirp using the Hilbert transform and use it to draw *Z*^+/-^(*f*). to obtain *Z*^+/-^(*f*) we interpolate the set {*Z*^+/-^(*f*)}.

### Sinusoidal stimulation at 4 Hz and spike-phase analysis

Cells were assigned to the L5 classes by clustering; when a genetic label was available, it was used instead of the cluster label. As above, a small holding current kept the resting membrane potential at about −70 mV during the whole recording. Cells received 10 cycles of a 4 Hz sinusoidal current. The amplitude was set to the cell’s cycle threshold, the lowest amplitude that evoked a spike on every cycle, adjusted in 20 pA increments; this differs from the step rheobase used for the AHP and adaptation measurements. Each cell received 2–7 repetitions, which were analysed separately and then averaged. The instantaneous phase of the injected current was obtained from its Hilbert transform, with 0° at the current peak and positive values after it. Spike phase was the phase of the current at the first spike of each cycle (measured relative to the spike peak). The first and last cycles were excluded to avoid onset and offset effects, leaving eight cycles (c0–c7). At 4 Hz, 1° corresponds to 0.69 ms.

For each cell, phases were unwrapped across cycles, and phase drift was the slope of a least-squares linear fit of phase against cycle number (°/cycle; negative values indicate an advance). Drift was tested against zero with a one-sample t-test across cells and reported with a t-based 95% confidence interval; per-cell slopes did not depart from normality (Shapiro–Wilk p = 0.12). Drift and per-cell mean phase (circular mean over c0–c7) were compared between classes with Kruskal–Wallis tests. In an exploratory analysis, drift in ET cells (ET1 and ET2 pooled) was tested against zero with an exact Wilcoxon signed-rank test. Cycle-to-cycle variation was the standard deviation of each cell’s unwrapped phase across c0–c7. For each cycle, the circular mean phase, the resultant length R and the circular standard error (circular SD/√n) were computed across cells. The cell was the unit of analysis throughout.

Analysis was done in Python 3.13 using NumPy 2.5 and SciPy 1.18 (code available upon request).

### In vivo electrophysiological recordings

Headplate implantation and the craniotomy were performed as previously described (Cano-Ferrer et al., 2024). Extracellular recordings were made from awake, head-fixed mice freely running on an air-supported Styrofoam ball. The medial secondary visual cortex (V2M) was targeted at 1.5 mm lateral to the midline and 1.2 mm anterior to the lambda suture. Recordings were made using linear silicon probes (A16-5mm-100-413; NeuroNexus), connected to a digitising headstage (PZ2-32) and acquired via an RZ2 processor (Tucker-Davis Technologies). For each animal, a single recording site in deep cortex near layer 5, selected by its depth along the probe, was used for analysis. Local field potentials (LFP) were extracted by filtering the raw signal between 1 and 100 Hz.

### In vivo visual stimulation

Visual stimuli were presented on a computer monitor to the eye contralateral to the recording hemisphere. Each trial comprised pre-and post-stimulus periods of uniform 50% grey for 10 seconds, flanking a 20 s stimulation epoch, during which the screen alternated between black and white (“flip” stimulus) at 1, 5 or 10 Hz (full black–white cycles per second).

### In vivo data analysis

Data extraction and analysis were done in Matlab. Time–frequency spectral content of the LFP was computed using a Morlet wavelet transform. Power spectra were computed by fast Fourier transform over 10 s windows from the pre-stimulus, stimulus and post-stimulus periods and are shown at 1 Hz resolution between 1 and 10 Hz. Spectra were normalised to the maximum pre-stimulus power and averaged across mice (n = 5); values are mean ± SEM.

## Results

### Subthreshold filtering separates IT and ET cells in V2M layer 5

We recorded from L5 cells in V2M in the presence of synaptic blockers to isolate intrinsic properties. To keep voltage-dependent conductances in a comparable state, a holding current kept each cell near −70 mV (−72 ± 0.37 mV, n = 51). A small (±20 pA) chirp current, increasing exponentially in frequency from 0.5 to 50 Hz (Fig. 1A), evoked approximately linear voltage responses, from which we computed the impedance profile (Methods), shown as the Nyquist plot of reactance against resistance (Fig. 1B) and as its magnitude (Fig. 1C) and phase (Fig. 1D). A non-resonant cell behaves like an RC circuit. It acts as a low-pass filter, and its voltage lags the current at all frequencies. A resonant cell behaves like an RLC circuit with a phenomenological inductance. Its impedance peaks at an intermediate frequency (the resonance frequency), and at low frequencies its voltage leads the current. We term the area of this positive-phase region the inductive phase, and the frequency at which the phase returns to zero, where the peaks of voltage and current coincide, the synchronous frequency (Fig. 1D).

**Figure 1.**
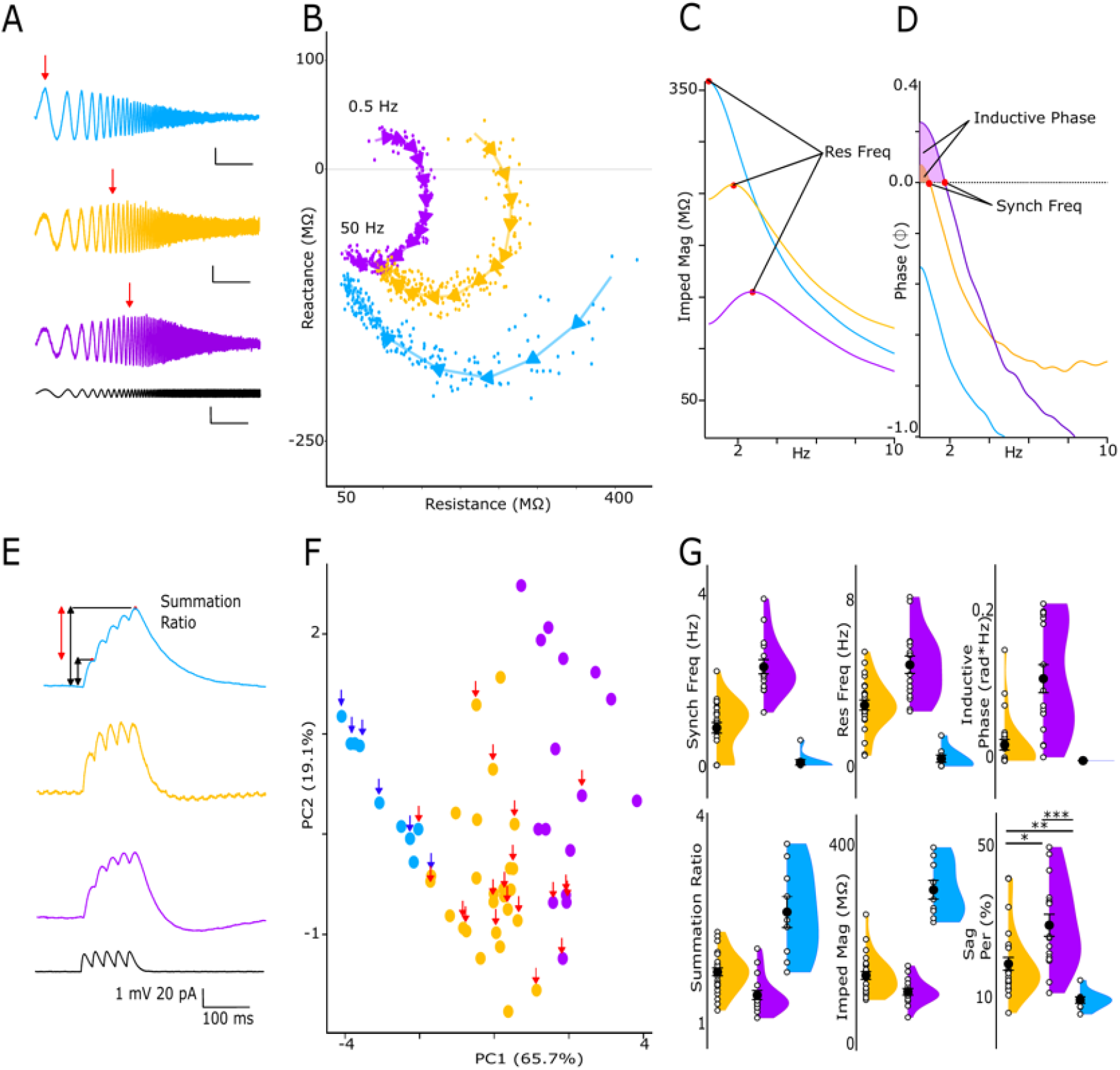
Subthreshold filtering separates IT and ET cells in V2M layer 5. **A.** Voltage responses of three cells from the IT (blue), ET1 (orange) and ET2 (purple) clusters to a ±20 pA chirp current (exponentially increasing from 0.5 to 50 Hz; bottom). Red arrows mark the resonance frequency. Scale bars: 5 mV (top), 4 mV (middle), 2 mV (bottom); 5 s. **B.** Nyquist plots (reactance against resistance) of the cells in A; points are frequency bins, and arrows show the direction of increasing frequency. **C.** Impedance magnitude profiles of the cells in A; red dots mark the resonance frequency. **D.** Impedance phase profiles of the cells in A; red dots mark the synchronous frequency, and shading above zero the inductive phase. **E.** Responses of the cells in A to five EPSP-like 20 pA current injections at 40 Hz. **F.** First two principal components of five intrinsic features (n = 51 cells; 84.8% of the variance), coloured by k-means cluster (IT, n = 9; ET1, n = 26; ET2, n = 16). Blue and red arrows mark Tlx3-Cre (IT) and Colgalt2-Cre (ET) cells. **G.** Resonance frequency, impedance magnitude, summation ratio, synchronous frequency, inductive phase and sag by cluster; mean ± SEM with individual cells. The first five are clustering features and are not compared between clusters; sag was compared with a Kruskal–Wallis test and Dunn’s post hoc tests with Bonferroni correction (* p < 0.05, ** p < 0.01, *** p < 0.001).

Ih, which contributes to resonance near rest (Hutcheon and Yarom, 2000), also reduces the temporal summation of EPSP trains (Magee, 1998, 1999; Remme and Rinzel, 2011; Mishra and Narayanan, 2025). We therefore also measured summation, as the ratio of the last to the first voltage peak in response to five EPSP-like 20 pA current injections at 40 Hz (Fig. 1E).

To relate these properties to projection class, we recorded from genetically labelled ET cells (Colgalt2-Cre::Ai14; n = 19) and IT cells (Tlx3-Cre::Ai14; n = 6) in addition to 26 cells from wild-type mice. For all 51 cells, we performed a principal component analysis on five features: resonance frequency, impedance magnitude, synchronous frequency, inductive phase and summation ratio (Fig. 1F). The first two principal components captured 84.8% of the variance, the first alone 65.7%. All five features contributed substantially to the first component: resonance frequency and impedance magnitude loaded most heavily (0.48 and −0.48), followed by synchronous frequency and summation ratio (0.47 and −0.47) and inductive phase (0.31) (Fig. S1A). The first component thus separated low-pass, strongly summating cells from resonant cells with lower input impedance.

We assessed the number of clusters for k-means clustering with silhouette scores and the within-cluster sum of squares (WCSS) (Fig. S1B, C). The WCSS curve declined without a clear elbow, and the silhouette score was highest for k = 2 (0.40) and lower and similar for k = 3–5 (0.34–0.37). Both the two-and the three-cluster solutions agreed with the genetic labels for 23 of 25 labelled cells; the three-cluster solution additionally divided the ET population into a moderate-resonance group (ET1) and a high-resonance group (ET2). We therefore treat ET1 and ET2 as resonance-defined subgroups of the ET population; whether they are discrete types or the ends of a continuum cannot be resolved with this dataset (see Discussion). Clusters are shown in blue (IT, n = 9), orange (ET1, n = 26) and purple (ET2, n = 16) in Fig. 1F. To assess how robustly each cell was assigned, we computed a stability score across k-means runs with different random seeds (Methods; Fig. S1D). The median stability score was 0.95, with 40 stable (core) and 11 borderline cells; borderline cells lay at the boundaries between groups.

IT cells summated EPSP-like inputs strongly (summation ratio 2.60 ± 0.22) and acted as low-pass filters: 7 of 9 were non-resonant, with maximal impedance at the lowest frequency tested, and 8 of 9 had no inductive phase or synchronous frequency. ET cells formed two groups: ET1, with reduced summation (1.73 ± 0.06) and moderate resonance frequency (2.93 ± 0.23 Hz), and ET2, with the weakest summation (1.40 ± 0.07) and the highest resonance frequency (4.87 ± 0.42 Hz) (Fig. 1G). ET1 cells had a small inductive phase (0.020 ± 0.006 rad·Hz) and, in the 19 of 26 cells whose phase crossed zero, a low synchronous frequency (1.18 ± 0.08 Hz); ET2 cells had a larger inductive phase and a higher synchronous frequency (0.10 ± 0.02 rad·Hz; 2.28 ± 0.17 Hz; all 16 cells).

The spike afterhyperpolarisation (AHP) and adaptation were measured from responses to a 500 ms step at twice rheobase (Fig. S2A). The AHP was larger in IT than in ET cells but did not differ between ET1 and ET2, and the adaptation ratio did not differ between clusters (Fig. S2C; Table 1). One cell had no AHP or adaptation values because its 2x rheobase trace was unanalysable.

Across cortical areas, including mouse visual cortex, ET cells show a larger hyperpolarisation-induced sag, a hallmark of Ih, than their IT neighbours (Kim et al., 2015). Consistent with this, sag in response to a 500 ms hyperpolarising step reaching about −100 mV (Fig. S2B), a measure not used for clustering, differed between clusters and followed the order ET2 > ET1 > IT (Fig. 1G; Table 1).

Inputs in vivo are often large enough to recruit voltage-dependent conductances, whose effects a linear impedance analysis does not capture (Kelley et al., 2023). We therefore examined how ET cells respond to large inputs.

### Large inputs reveal asymmetric gain in ET cells

To characterise how ET cells respond to large inputs, we injected a ±80 pA chirp current, increasing linearly in frequency from 0.2 to 20 Hz, into the soma in the presence of synaptic blockers (Fig. 2A). The subthreshold response was non-linear: depolarising peaks and hyperpolarising troughs differed, consistent with voltage-dependent conductances acting on either side of the holding potential. Following Pena et al. (2019) and Kelley et al. (2023), we extracted impedance magnitude and phase profiles separately for the peaks and the troughs (Fig. 2B; Methods). This analysis was done in nine wild-type cells classified as ET by clustering, ET1 (n = 2) or ET2 (n = 7) (Fig. 1F); none was genetically or projection-identified. Resonance frequency, from the magnitude profiles, did not differ between troughs and peaks (3.61 ± 0.25 vs 3.16 ± 0.26 Hz; Fig. 2C; Table 1). Impedance magnitude was larger for peaks than for troughs (125.6 ± 12.4 vs 110.9 ± 11.4 MΩ; Fig. 2C; Table 1), by 14 ± 4% on average and in 8 of 9 cells. In the phase profiles, troughs tended to have a larger inductive phase than peaks (0.14 ± 0.03 vs 0.05 ± 0.03 rad·Hz; Fig. 2D). The peak response had no inductive phase, and hence no synchronous frequency, in 4 of 9 cells, and the trough response in 1 of 9. Of the four measures, only impedance magnitude differed significantly after Bonferroni correction (Table 1). The pattern is consistent with predictions from models of L5 pyramidal-tract neurons, in which depolarising half-cycles show less phase lead and a lower synchronous frequency than hyperpolarising ones, resonance frequency is nearly unchanged and the depolarising response is slightly larger (Kelley et al., 2023). The larger gain for depolarising half-cycles is consistent with Ih, which deactivates on depolarisation and activates on hyperpolarisation; in models of L5 pyramidal-tract neurons, blocking Ih abolishes the difference in phase between depolarising and hyperpolarising half-cycles (Kelley et al., 2023).

**Figure 2.**
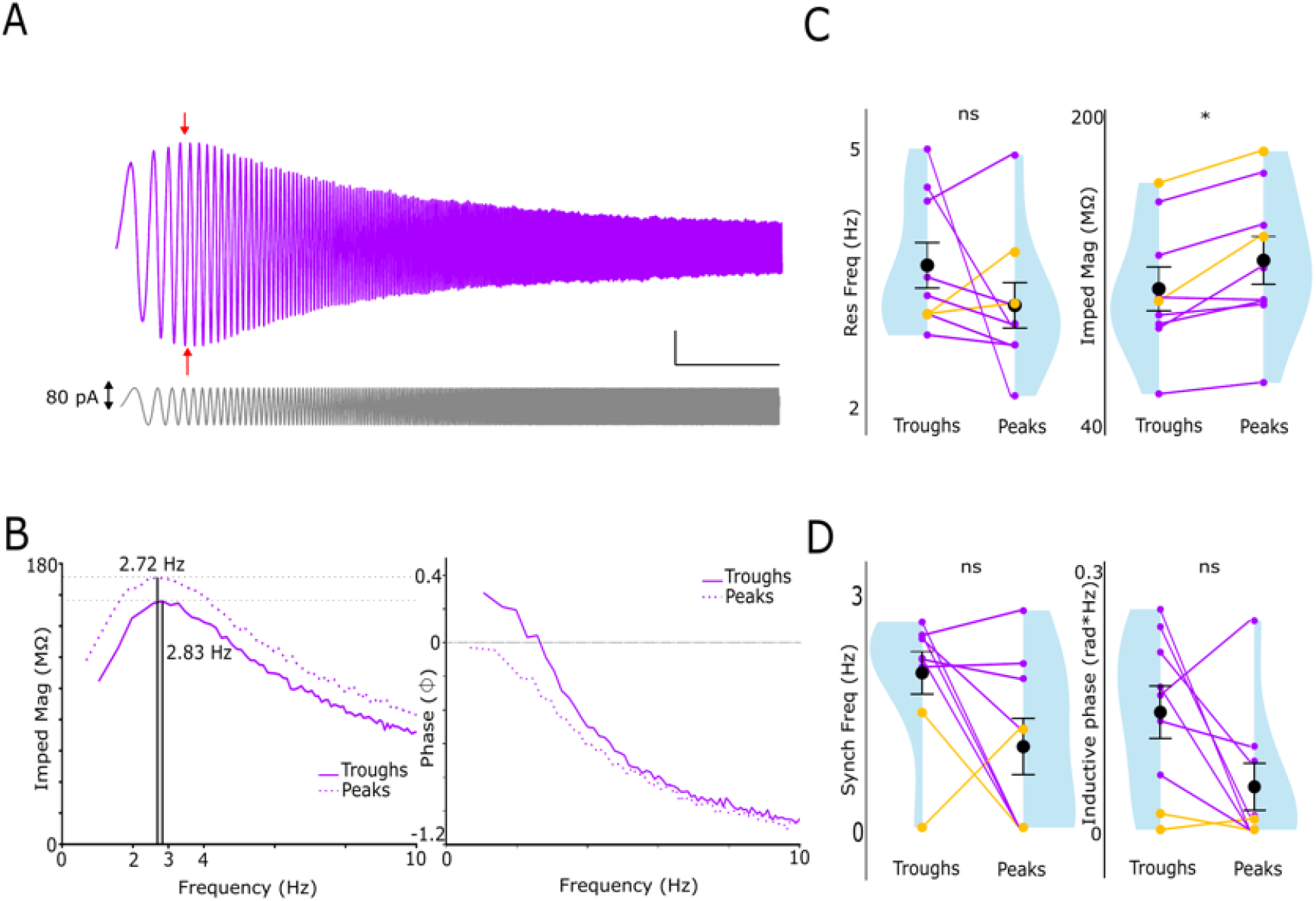
Large inputs reveal asymmetric gain in ET cells. **A.** Voltage response (top) of an ET2 cell to a large chirp current (±80 pA, increasing linearly from 0.2 to 20 Hz; bottom). Red arrows mark the resonance frequencies of the peaks and troughs. Scale bars: 5 mV, 5 s. **B.** Impedance magnitude (left) and phase (right) profiles of the cell in A, computed separately for troughs (solid) and peaks (dotted). Vertical lines mark the resonance frequencies of troughs (2.83 Hz) and peaks (2.72 Hz); horizontal dotted lines, their impedance maxima. **C.** Resonance frequency and impedance magnitude of troughs and peaks (n = 9 wild-type cells: ET1, orange, n = 2; ET2, purple, n = 7). **D.** Synchronous frequency and inductive phase of troughs and peaks (same cells). Synchronous frequency is undefined when the phase does not cross zero (plotted as 0). C, D: mean ± SEM with individual cells; paired t-tests, Bonferroni-corrected across the four measures (* p < 0.05).

### Spike phase changes little across cycles of sustained 4 Hz drive in all L5 classes

The L5 classes differ in how they filter small inputs (Fig. 1), and ET cells respond asymmetrically to large ones (Fig. 2). We next asked whether these differences extend to spike timing when the input is strong enough to evoke a spike on every cycle, and whether spike timing remains stable over a sustained bout of input. Spike timing under sustained rhythmic input need not be stationary: in models of L5 pyramidal-tract neurons, spike phase can drift across successive cycles of sinusoidal current, later or earlier depending on the frequency and amplitude of the input (Kelley et al., 2023). We injected 10 cycles of 4 Hz sinusoidal current into IT (n = 6), ET1 (n = 5) and ET2 (n = 7) cells in the presence of synaptic blockers (Fig. 3A); ET1 and ET2 cells were wild-type cells classified by clustering, and IT cells were five Tlx3-Cre cells and one wild-type cell of the IT cluster (genetic identity overrides cluster identity). The amplitude was set to each cell’s cycle threshold, the lowest amplitude that evoked a spike on every cycle, so that spike timing was compared at matched drive relative to threshold. Spike phase was measured at the first spike of each cycle, relative to the injected current (0° = current peak; positive values = after the peak). The first and last cycles were excluded, leaving eight cycles (c0–c7; Methods).

**Figure 3.**
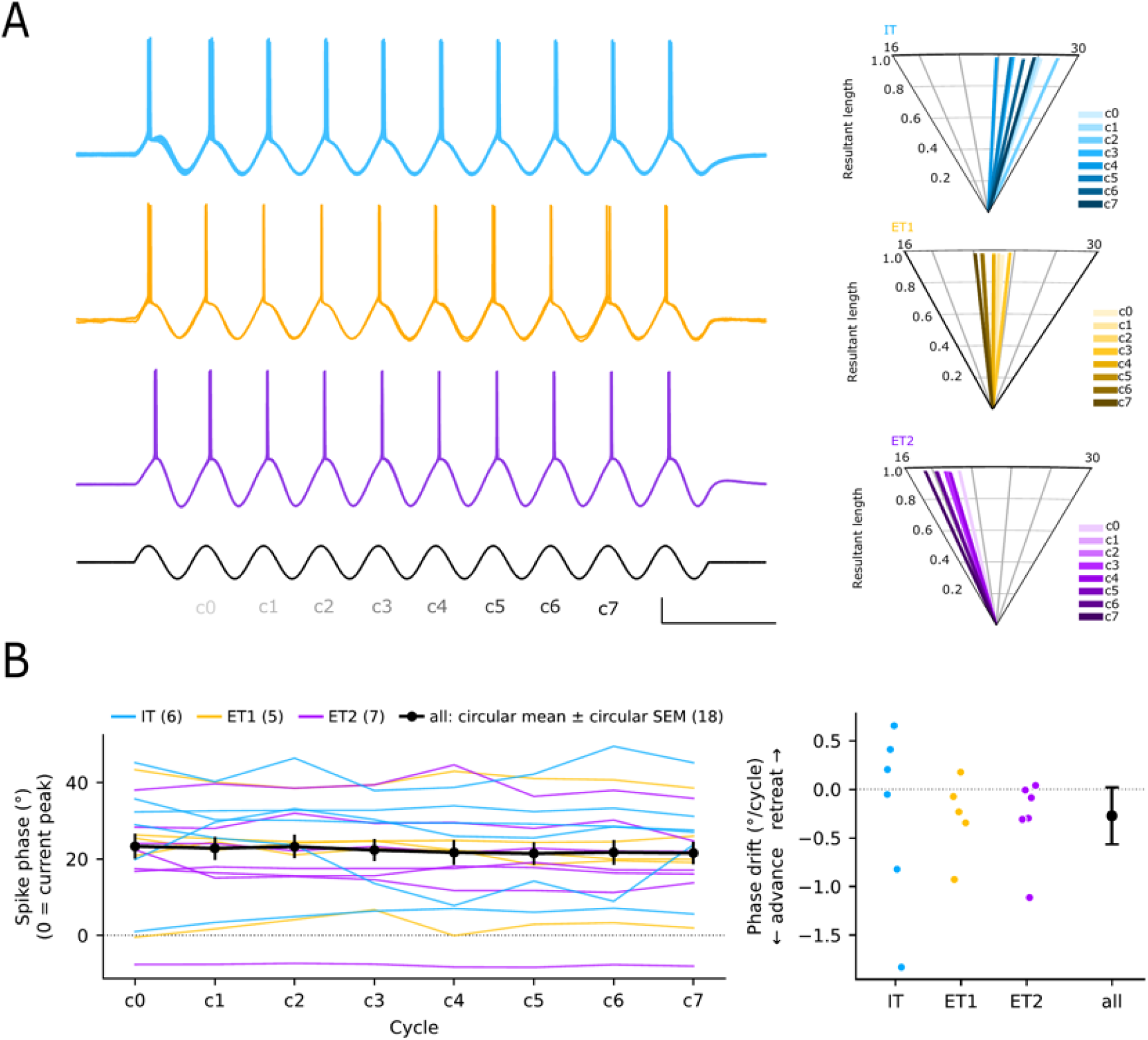
Spike phase changes little across cycles of 4 Hz drive in all L5 classes. **A.** Voltage responses of an IT (blue), an ET1 (orange) and an ET2 (purple) cell to 10 cycles of 4 Hz sinusoidal current (bottom) at the cell’s threshold amplitude, the lowest amplitude that evoked a spike on every cycle. Scale bars: 10 mV, 120 pA. The first and last cycles were excluded from the phase analysis; the remaining cycles are labelled c0–c7. Right, for each class: circular mean spike phase (angle; 16–30°, angular axis expanded) and resultant length R (radius) at each cycle c0–c7 (light to dark), across the cells of that class. **B.** (Left) Spike phase across cycles c0–c7: the phase of the injected current at the first spike of each cycle (0° = current peak; positive = after the peak). Thin lines, individual cells coloured by class (IT n = 6, ET1 n = 5, ET2 n = 7); black, circular mean ± circular SEM across all cells (n = 18). The resultant length was 0.97–0.98 at every cycle. (Right) Phase drift of each cell: slope of a linear fit of its unwrapped spike phase against cycle (c0-c7). Dots, individual cells by class; black, mean and 95% CI across all cells. Slopes did not differ between classes. IT: Tlx3-Cre cells and one wild-type cell of the IT cluster; ET1 and ET2: wild-type cells assigned by clustering (Fig. 1).

Spikes occurred near the peak of the injected current (Fig. 3A, B): the mean spike phase across cells was 22 ± 3° (circular mean ± circular SEM; about 15 ms after the peak), with per-cell means from −8° to 43°, and at every cycle the resultant length across cells was 0.97–0.98, so cells fired at similar phases. Mean spike phase did not differ between classes (IT 26 ± 5°, ET1 22 ± 5°, ET2 19 ± 5°; Table 1).

Spike phase changed little across cycles (Fig. 3B). Each cell’s phase varied only slightly from cycle to cycle (SD across the eight cycles: median 2.1°, interquartile range 1.0–2.7°) compared with the spread of mean phase between cells (circular SD 13°). A linear fit of each cell’s phase against cycle gave a mean drift of −0.27 ± 0.14°/cycle (0.19 ms per cycle; 95% CI −0.57 to +0.02; 13 of 18 cells advanced; Fig. 3B, right), which did not differ significantly from zero (Table 1); the upper confidence limit excludes a retreat of more than 0.02°/cycle. Drift did not differ between classes (Fig. 3B, right; Table 1). In an exploratory analysis of ET cells (ET1 and ET2 pooled), drift was significantly negative (10 of 12 cells advanced; median −0.26°/cycle, 0.18 ms per cycle; Table 1), whereas IT cells showed no consistent drift (3 of 6 advanced). Because drift did not differ between classes, the advance cannot be attributed specifically to ET cells.

### Visual flip stimulation increases 1–6 Hz LFP power in deep layers of V2M

In vitro, ET cells in V2M resonated at about 3 Hz (ET1) and 5 Hz (ET2), whereas IT cells did not, and when driven to fire at 4 Hz, all classes kept a nearly constant spike phase. In awake mice, brief visual stimuli evoke bouts of 5 Hz oscillation in the pyramidal cell population of V1 and retrosplenial cortex (White et al., 2026). Whether rhythmic activity in this range also reaches the deep layers of V2M, where ET and IT cells reside, has not been tested. To address this, we recorded LFPs with linear silicon probes in V2M of awake, head-fixed mice and presented full-screen flip stimuli to the contralateral eye, analysing a recording site in deep cortex near layer 5. A 1 Hz flip stimulus evoked prominent oscillation bouts faster than the stimulus (Fig. 4A). In the group-averaged spectra, LFP power during stimulation was higher than before or after it across about 1–6 Hz (Fig. 4B; 5 mice, 39 sessions).

**Figure 4.**
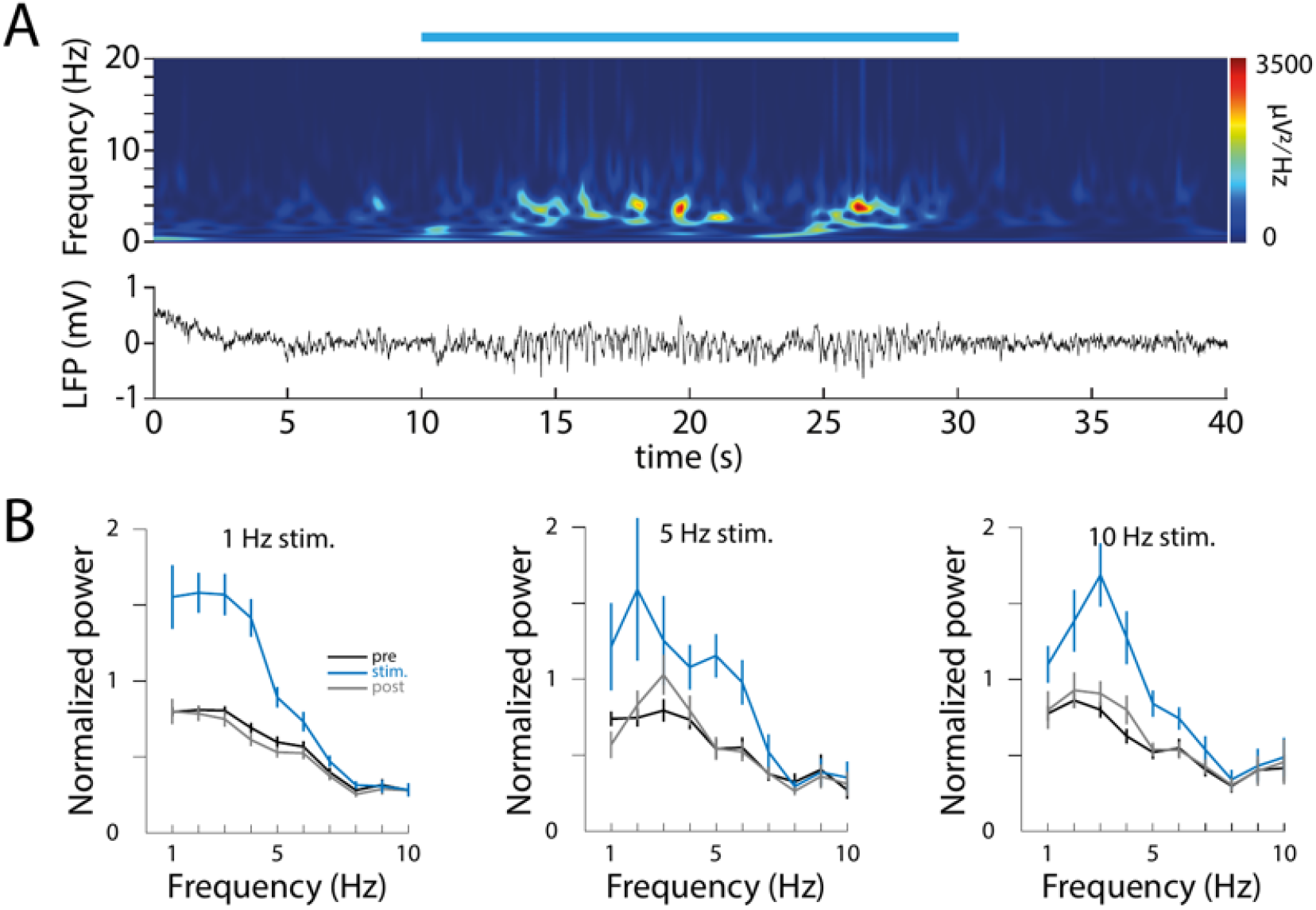
Visual flip stimulation increases 1–6 Hz LFP power in deep layers of V2M, spanning the ET resonance frequencies. **A.** Example recording from V2M with a 1 Hz full-screen flip stimulus (blue bar, 20 s). During stimulation, oscillations faster than the stimulus are visible in the LFP (bottom) and its Morlet wavelet spectrogram (top). **B.** LFP power spectra (1–10 Hz) before (black), during (blue) and after (grey) full-screen flip stimulation at 1 Hz (39 sessions), 5 Hz (11 sessions) and 10 Hz (13 sessions); 5 mice. Spectra were normalised to the maximum pre-stimulus power and averaged across mice; mean ± SEM (n = 5 mice).

At 1 Hz, however, responses locked to the luminance reversals and their harmonics also fall within this band. We therefore repeated the stimulation at 5 and 10 Hz (full black–white cycles per second). The increase across about 1–6 Hz was again present, peaking at 2–3 Hz (5 Hz: 11 sessions; 10 Hz: 13 sessions; 5 mice; Fig. 4B). In contrast, power at the drive frequency itself was similar before, during and after 10 Hz stimulation. As the band did not follow the drive frequency, it is unlikely to reflect stimulus-locked responses, and it is consistent with an oscillation that the stimulus triggers but does not pace. This band spans the mean resonance frequencies of ET1 and ET2 cells measured in vitro.

## Discussion

We recorded from layer 5 projection neurons in V2M to determine how their intrinsic properties shape responses to slow rhythmic input in three stimulus regimes, and tested whether a rhythm in this frequency range occurs in V2M in vivo. IT and ET neurons differed in their subthreshold filtering: IT neurons summated strongly and acted as low-pass filters, whereas ET neurons resonated, forming moderate-resonance (ET1) and high-resonance (ET2) subgroups. With large inputs, ET neurons amplified depolarisation more than hyperpolarisation. When driven to fire at 4 Hz, neurons of all classes fired at a similar phase of the input and kept it from cycle to cycle. In awake mice, visual stimulation increased deep-layer LFP power across about 1–6 Hz, a band that spans the mean resonance frequencies of ET1 and ET2 neurons.

The division between IT and ET neurons was evident in both temporal summation and resonance. Our results extend to a higher-order visual area of the mouse a dissociation first described in rat medial prefrontal cortex, where ET neurons act as band-pass filters in the theta range and summate weakly, whereas IT neurons act as low-pass filters and summate strongly (Dembrow et al., 2010). Within the ET population, clustering separated a moderate-resonance group (mean about 3 Hz) from a high-resonance group (mean about 5 Hz), although their resonance ranges overlapped. Sag followed the same order (ET2 > ET1 > IT), consistent with graded Ih. However, as sag, resonance and summation all depend on Ih, this agreement supports the ordering of the groups but does not provide independent evidence that they are distinct.

With large inputs, the responses of ET neurons became asymmetric. Resonance frequency was similar for depolarising peaks and hyperpolarising troughs (about 3–4 Hz), but peaks were amplified more than troughs. Troughs also tended to lead the current more than peaks, although this difference was not significant after correction (corrected p = 0.20). This pattern is consistent with modelling of L5 pyramidal-tract neurons, which predicted that depolarising and hyperpolarising half-cycles are processed with different phase once inputs are large enough to engage voltage-dependent conductances asymmetrically, with little change in resonance frequency (Kelley et al., 2023). The gain asymmetry we measured (about 14%) was, however, larger than in those simulations (about 5%). The lower gain of the troughs, and their tendency to lead more, are consistent with Ih acting as a restoring current on the hyperpolarised side (Mishra and Narayanan, 2025). In contrast, the larger gain of the peaks argues against a dominant M-current on the depolarised side, since an outward, depolarisation-activated current would reduce rather than increase depolarising gain. A more likely explanation is that depolarisation deactivates Ih and thereby raises membrane resistance, possibly with a contribution from amplifying inward currents such as the persistent sodium current (Hutcheon and Yarom, 2000). The M-current may contribute a small phase lead at the peaks but is unlikely to dominate in this voltage range. In the models, blocking Ih abolishes the phase difference between half-cycles (Kelley et al., 2023). As we did not block any of these conductances, this account is inferred from the impedance profiles; applying ZD7288 (Ih blocker), XE991 (M-current blocker) or low-dose TTX (sodium channel blocker) would distinguish between these contributions.

Once driven to fire, the classes did not differ detectably. At 4 Hz, with drive set to each cell’s threshold, spikes in all classes occurred near the current peak and kept their phase from cycle to cycle. The mean drift was an advance of about 0.2 ms per cycle (95% CI from a 0.4 ms advance to a 0.01 ms delay per cycle). This suggests that adaptation building up across cycles is weak in these neurons under these conditions. In models of L5 pyramidal-tract neurons, the phase of spiking under sustained sinusoidal drive can retreat or advance across cycles, depending on the input frequency and on how its amplitude changes (Kelley et al., 2023). As we tested a single frequency at threshold drive, it remains to be seen whether V2M neurons shift their spike phase under other conditions. Under these conditions, spike phase thus provides a near-constant reference. This stability should be distinguished from hippocampal phase precession, which unfolds over many cycles and reflects the interaction of a cell’s firing with a network rhythm during behaviour; here, we examined single neurons under fixed drive.

Widefield voltage imaging has shown that brief visual stimuli evoke bouts of 5 Hz oscillation in V1 and retrosplenial cortex (White et al., 2026). We found a related but broader increase in the deep layers of V2M, where visual flip stimulation increased LFP power across about 1–6 Hz, peaking at 2–3 Hz. As a square-wave flip stimulus contains harmonics, the 1 Hz condition alone could account for elevated power at low multiples of the drive frequency. Responses to flicker can also show period doubling, a component at half the stimulus frequency (Crevier and Meister, 1998), which at 5 Hz would fall at 2.5 Hz. The 10 Hz condition is therefore the most informative. In the group-averaged spectra, power was again higher across about 1–6 Hz, peaking at 2–3 Hz, below both the drive frequency and half of it, while power at 10 Hz itself was similar before, during and after stimulation. The band thus did not follow the stimulus, which is consistent with an oscillation that the stimulus triggers but does not pace. Similarly, gratings lasting 0.2–0.5 s evoke 5 Hz bouts lasting about 0.7 s in V1 (White et al., 2026). Widefield glutamate imaging in mouse visual cortex likewise showed more 3–5 Hz power during flicker than between trials, more so for flicker below 20 Hz, and not confined to visual areas; the authors tentatively related it to breathing-linked 4 Hz oscillations (Gulbinaite et al., 2024). The origin of this rhythm remains open. The co-occurrence of visually evoked 5 Hz oscillations in V1 and retrosplenial cortex (White et al., 2026), together with the retrosplenial input to V2M ET neurons (Galloni et al., 2022), makes a top-down source plausible. On the other hand, paired recordings across visual thalamus, retrosplenial cortex and hippocampus suggest that familiarity-evoked theta (4–8 Hz) in V1 is generated within visual cortex, with a delayed onset in retrosplenial cortex (Zimmerman et al., 2025). In either case, ET neurons are unlikely to pace the rhythm, as they form the output of the cortical column and provide little local excitatory output (Harris and Shepherd, 2015). Their intrinsic properties instead suit them to read out the rhythm.

Several caveats should be considered. The division of ET neurons into ET1 and ET2 rests on clustering of resonance and summation features and is mirrored by sag, all of which depend on Ih. Whether these groups represent discrete subtypes or the ends of a graded distribution is not resolved by our data, and there is at present no transcriptomic, morphological or projection-based evidence for such a division within the Colgalt2-Cre population. We therefore use the group labels descriptively. The stimuli were delivered as somatic current injection, which does not reproduce the spatial distribution of synaptic input. However, retrosplenial input to V2M ET neurons peaks on perisomatic dendrites in most neurons (Galloni et al., 2022), so somatic integration is relevant for this pathway. Recordings in slices also lack the tonic synaptic bombardment of the intact cortex, which alters membrane conductance and could shift resonance. The large-input analysis was also restricted to ET neurons, so we cannot say whether IT neurons show the same asymmetry. Finally, the in vivo results are descriptive group averages from five mice, and the recordings measure the LFP without unit isolation or cell-type identification; the relationship between the intrinsic resonance of ET neurons and the evoked band is a correspondence in frequency, not a demonstration that these cells take part in it.

In summary, the two output channels of layer 5 in V2M, which carry information to different targets, filter slow rhythmic input differently. IT neurons act as low-pass filters without a preference for frequencies in this range, whereas ET neurons resonate at frequencies within the band evoked by visual stimulation in vivo. Once driven to fire, however, both keep a nearly constant spike phase. A bout of such rhythmic input arriving in layer 5 would therefore be filtered differently by its two output channels but, as far as we could detect, timed alike.

## Data and code availability

Data and analysis code are available from the corresponding authors on request.

## Funding

This work was funded (E.R.) by the Excellence Initiative of Aix-Marseille Université, A*Midex (AMX-20-CE-06), a French “Investissements d’Avenir” programme. Further support (to H.T.) was received from the French government under the “France 2030” programme via A*Midex (AMX-19-IET-004) and ANR (ANR-17-EURE-0029) funding, as well as Wellcome Trust (to E.R., 104285/B/14/Z) and by the Francis Crick Institute (to E.R. and Z.Y.) which receives its core funding from Cancer Research UK (FC001153), the UK Medical Research Council (FC001153), and the Wellcome Trust (FC001153).

## Author contributions

HT: Conceptualization, Formal analysis, Investigation, Writing - original draft, Writing - review and editing; ZY: Formal analysis, Investigation; ER: Conceptualization, Supervision, Funding acquisition, Methodology, Writing - original draft, Project administration, Writing - review and editing

## Acknowledgements

We thank Thomas Vannier for help with PCA analysis and clustering; Thais Barbosa Caetano Souza and Rishikesh Narayanan for advice on analysis; Laurent Aniksztejn for advice on experiments; and all members of the lab for advice and support.

## Conflict of interest

None

---

**Figure S1.**
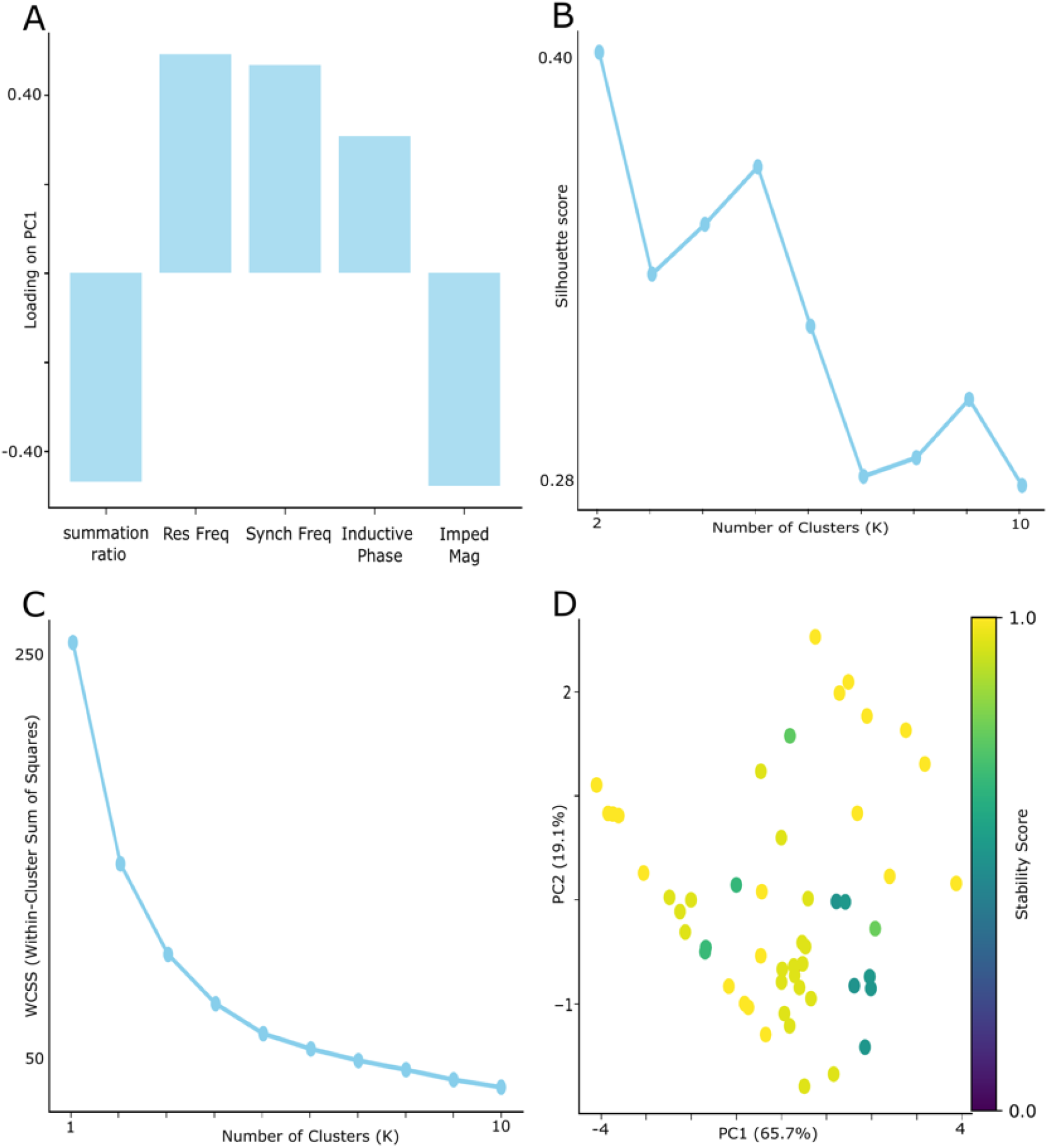
Principal component analysis and choice of cluster number. **A.** Loading of each feature on the first principal component. **B.** Mean silhouette score for each number of clusters. **C.** Within-cluster sum of squares for each number of clusters (elbow plot). **D.** Principal component plot as in Fig. 1F, with each cell coloured by its stability score across k-means runs with different random seeds (k = 3; Methods).

**Figure S2.**
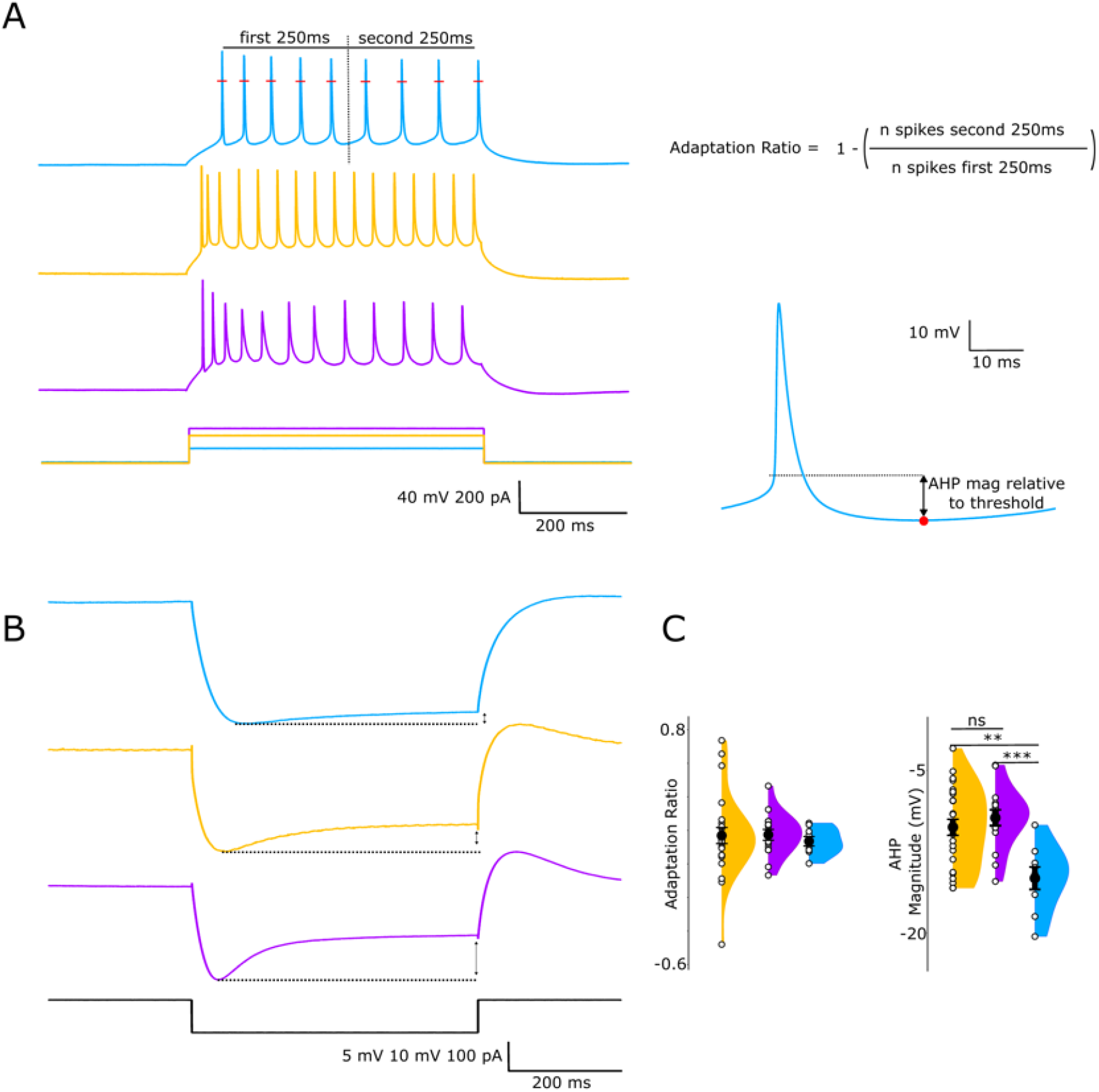
Afterhyperpolarisation, adaptation and sag. **A.** Responses of the example cells in Fig. 1A to a 500 ms depolarising step at twice rheobase. The adaptation ratio compares the number of spikes in the first and second halves of the step; the AHP was measured relative to spike threshold on the average spike waveform. **B.** Responses of the same cells to a 500 ms hyperpolarising step reaching about −100 mV, used to measure sag. **C.** Adaptation ratio and AHP by cluster; mean ± SEM with individual cells (IT n = 9, ET1 n = 25, ET2 n = 16). AHP: one-way ANOVA with Tukey’s post hoc tests; adaptation: Kruskal–Wallis test (** p < 0.01, *** p < 0.001).

## Notes

### Competing Interest Statement

The authors have declared no competing interest.

## References

Buzsáki, G., 2002. Theta Oscillations in the Hippocampus. Neuron, 33(3), pp.325–340. doi:10.1016/s0896-6273(02)00586-x

Cano-Ferrer, X., Tran-Van-Minh, A. and Rancz, E., 2024. RPM: An open-source Rotation Platform for open-and closed-loop vestibular stimulation in head-fixed Mice. Journal of Neuroscience Methods, 401, 110002. doi:10.1016/j.jneumeth.2023.110002

Colgin, L.L., 2013. Mechanisms and Functions of Theta Rhythms. Annual Review of Neuroscience, 36, pp.295–312. doi:10.1146/annurev-neuro-062012-170330

Crevier, D.W. and Meister, M., 1998. Synchronous Period-Doubling in Flicker Vision of Salamander and Man. Journal of Neurophysiology, 79(4), pp.1869–1878. doi:10.1152/jn.1998.79.4.1869

Dembrow, N.C., Chitwood, R.A. and Johnston, D., 2010. Projection-Specific Neuromodulation of Medial Prefrontal Cortex Neurons. The Journal of Neuroscience, 30(50), pp.16922–16937. doi:10.1523/JNEUROSCI.3644-10.2010

Dura-Bernal, S., Neymotin, S.A., Suter, B.A., Dacre, J., Moreira, J.V.S., Urdapilleta, E., Schiemann, J., Duguid, I., Shepherd, G.M.G. and Lytton, W.W., 2023. Multiscale model of primary motor cortex circuits predicts in vivo cell-type-specific, behavioral state-dependent dynamics. Cell Reports, 42(6), 112574. doi:10.1016/j.celrep.2023.112574

Fournier, J., Saleem, A.B., Diamanti, E.M., Wells, M.J., Harris, K.D. and Carandini, M., 2020. Mouse Visual Cortex Is Modulated by Distance Traveled and by Theta Oscillations. Current Biology, 30(19), pp.3811–3817.e6. doi:10.1016/j.cub.2020.07.006

Galloni, A.R., Ye, Z. and Rancz, E., 2022. Dendritic Domain-Specific Sampling of Long-Range Axons Shapes Feedforward and Feedback Connectivity of L5 Neurons. The Journal of Neuroscience, 42(16), pp.3394–3405. doi:10.1523/JNEUROSCI.1620-21.2022

Gulbinaite, R., Nazari, M., Rule, M.E., Bermudez-Contreras, E.J., Cohen, M.X., Mohajerani, M.H. and Heimel, J.A., 2024. Spatiotemporal resonance in mouse primary visual cortex. Current Biology, 34(18), pp.4184–4196.e7. doi:10.1016/j.cub.2024.07.091

Harris, K.D. and Shepherd, G.M., 2015. The neocortical circuit: themes and variations. Nature Neuroscience, 18(2), pp.170–181. doi:10.1038/nn.3917

Hutcheon, B. and Yarom, Y., 2000. Resonance, oscillation and the intrinsic frequency preferences of neurons. Trends in Neurosciences, 23(5), pp.216–222. doi:10.1016/s0166-2236(00)01547-2

Ito, J., Roy, S., Liu, Y., Cao, Y., Fletcher, M., Lu, L., Boughter, J.D., Grün, S. and Heck, D.H., 2014. Whisker barrel cortex delta oscillations and gamma power in the awake mouse are linked to respiration. Nature Communications, 5, 3572. doi:10.1038/ncomms4572

Kelley, C., Dura-Bernal, S., Neymotin, S.A., Antic, S.D., Carnevale, N.T., Migliore, M. and Lytton, W.W., 2021. Effects of Ih and TASK-like shunting current on dendritic impedance in layer 5 pyramidal-tract neurons. Journal of Neurophysiology, 125(4), pp.1501–1516. doi:10.1152/jn.00015.2021

Kelley, C., Antic, S.D., Carnevale, N.T., Kubie, J.L. and Lytton, W.W., 2023. Simulations predict differing phase responses to excitation vs. inhibition in theta-resonant pyramidal neurons. Journal of Neurophysiology, 130(4), pp.910–924. doi:10.1152/jn.00160.2023

Kim, E.J., Juavinett, A.L., Kyubwa, E.M., Jacobs, M.W. and Callaway, E.M., 2015. Three Types of Cortical Layer 5 Neurons That Differ in Brain-wide Connectivity and Function. Neuron, 88(6), pp.1253–1267. doi:10.1016/j.neuron.2015.11.002

Lisman, J.E. and Jensen, O., 2013. The Theta-Gamma Neural Code. Neuron, 77(6), pp.1002–1016. doi:10.1016/j.neuron.2013.03.007

Lur, G., Vinck, M.A., Tang, L., Cardin, J.A. and Higley, M.J., 2016. Projection-Specific Visual Feature Encoding by Layer 5 Cortical Subnetworks. Cell Reports, 14(11), pp.2538–2545. doi:10.1016/j.celrep.2016.02.050

Magee, J.C., 1998. Dendritic Hyperpolarization-Activated Currents Modify the Integrative Properties of Hippocampal CA1 Pyramidal Neurons. The Journal of Neuroscience, 18(19), pp.7613–7624. doi:10.1523/JNEUROSCI.18-19-07613.1998

Magee, J.C., 1999. Dendritic Ih normalizes temporal summation in hippocampal CA1 neurons. Nature Neuroscience, 2(6), pp.508–514. doi:10.1038/9158

Mishra, P. and Narayanan, R., 2025. The enigmatic HCN channels: A cellular neurophysiology perspective. Proteins: Structure, Function, and Bioinformatics, 93(1), pp.72–92. doi:10.1002/prot.26643

Pena, R.F.O., Lima, V., Shimoura, R.O., Ceballos, C.C., Rotstein, H.G. and Roque, A.C., 2019. Asymmetrical voltage response in resonant neurons shaped by nonlinearities. Chaos: An Interdisciplinary Journal of Nonlinear Science, 29(10), 103135. doi:10.1063/1.5110033

Remme, M.W. and Rinzel, J., 2011. Role of active dendritic conductances in subthreshold input integration. Journal of Computational Neuroscience, 31(1), pp.13–30. doi:10.1007/s10827-010-0295-7

Schroeder, C.E. and Lakatos, P., 2009. Low-frequency neuronal oscillations as instruments of sensory selection. Trends in Neurosciences, 32(1), pp.9–18. doi:10.1016/j.tins.2008.09.012

Tang, L. and Higley, M.J., 2020. Layer 5 Circuits in V1 Differentially Control Visuomotor Behavior. Neuron, 105(2), pp.346–354.e5. doi:10.1016/j.neuron.2019.10.014

White, C.M., Azimi, Z., Staadt, R., Song, C., Knöpfel, T. and Jancke, D., 2026. Psychedelic 5-HT2A agonist increases spontaneous and evoked 5-Hz oscillations in visual and retrosplenial cortex. Communications Biology, 9(1), 216. doi:10.1038/s42003-025-09492-9

Zimmerman, M.P., Kissinger, S.T., Edens, P., Towers, R.C., Nareddula, S., Nadew, Y.Y., Quinn, C.J. and Chubykin, A.A., 2025. Origin of visual experience-dependent theta oscillations. Current Biology, 35(1), pp.87–99.e6. doi:10.1016/j.cub.2024.11.015

